# New method for runoff estimation under different soil management practices

**DOI:** 10.1101/424069

**Authors:** Janvier Bigabwa Bashagakule, Vincent Logah, Andrews Opoku, Henry Oppong Tuffour, Joseph Sarkodie-Addo, Charles Quansah

## Abstract

Soil erosion has been widely measured using different approaches based on models, direct runoff and sediment collections. However, most of the methods are, poorly applied due to the cost, the accuracy and their tedious nature. This study aimed to develop and test a new method for runoff characterization, which may be more applicable and adaptable to different situations of soil and crop management. An experiment was carried out on runoff plots under different cropping systems (sole maize, sole soybean and maize intercropped with soybean) and soil amendments (NPK, Biochar, NPK + Biochar and Control) in the Semi-deciduous forest zone of Ghana. The study was a two-factor experiment (split-plot) in which cropping systems constituted the main plot whereas soil management the subplot. To assess the quality of the method, different statistical parameters were used: p-values, coefficient of determination (R^2^), Nash-Sutcliffe efficiency (NSE), root mean square (RMSE) and, root square ratio (RSR). The NPK + Biochar under each cropping system reduced surface runoff than all other treatments. At p < 0.001, R^2^ ranged from 0.88 to 0.94 which showed good accuracy of the method developed. The dispersion between the predicted and observed values was low with RMSE varying from 1.68 to 2.66 mm which was less than 10 % of the general mean of the runoff. Moreover, the low variability between parameters was confirmed by the low values of RSR ranging from 0.38 to 0.46 (with 0.00 ≤ RSR ≤ 0.50 for perfect prediction). NSE values varied from 0.79 to 0.86 (≥0.75 being the threshold for excellent prediction). Though the sensitivity analysis showed that the method under high amount of runoff (especially on bare plots) was poorly adapted, the dimensions of runoff plots could be based on runoff coefficient of the region by analyzing the possible limits of an individual rainfall amount of the site. The findings provide alternative approach for monitoring soil degradation.

## 1. Introduction

Characterizing soil erosion on the field is a critical option to sustain crop productivity due to its effect on the environment and on crop development [1]. Describing and quantifying the rate of soil erosion in a watershed over spatial and time scales is one of the constraints to direct soil erosion assessment due to the limits in field measurement [2] and the significant amount of sediments and runoff to handle. Adapted interventions are therefore clearly required to investigate the effect of climate and land use change, as the driver of rainwater fate on erosion rates towards the recommendation of sustainable land management practices.

Due to the constraints to the direct soil loss quantification, different and specific models and equations have been widely used to predict soil erosion over a wide range of conditions [3,4, 5, 6]. Most of the developed models are site-based equations making them more applicable to specific agro-ecological conditions [7] without a general adaptation to different ecosystems. They vary significantly in terms of their capability and complexity, input requirement, representation of processes, spatial and temporal scale accountability, practical applicability and with the types of output they provide [2]. For the applicability, each desirable model of soil erosion rate assessment should satisfy specific conditions of universal acceptability; reliability; robustness in nature; ease of use with minimum data set; and ability to take account of changes in land use, climate and conservation practices [2] . Apart from modeling by prediction, direct soil erosion measurement involves the use of big containers to harvest runoff but with poor success [8,9].

The use of automatic tipping buckets is one of the options for direct quantification of soil runoff and sediments with good accuracy[10] . However, the different stakeholders involved in soil and water conservation practices perceive this method, as very tedious and costly for adoption. Indeed, soil erosion measurement using direct and indirect approaches have been challenging in different studies due to the accuracy of the method and the important parameters required [11, 12]. Due to the various constraints to the tipping bucket and other methods of soil erosion characterization, there is a need to develop more useful and adapted approach based on numerical method which provides new options of assessing accurately soil runoff. This study therefore aimed to develop and test a new method to measure surface runoff on the field to reduce the constraints related to direct and indirect measurements. Moreover, it aimed to evaluate cropping systems and soil amendment combinations that can most effectively reduce runoff generated under rainfed cropping conditions in sub-Saharan (SSA).

## 2. Materials and methods

### 2.1. Research area description and field layout

The field experiment was carried out at the Anwomaso Agricultural Research Station of the Kwame Nkrumah University of Science and Technology, Kumasi, Ghana. The site is located within the semi-deciduous forest zone of Ghana and lies on longitude 1.52581° W and latitude 6.69756° N. The zone is characterized by two cropping seasons: March to July as the major season and September to December being the minor season. The rainfall pattern therefore is bimodal, ranging between 1300 and 1500 mm.

Runoff plots were installed based on two factors: cropping systems (Maize + soybean intercrop, sole maize, sole soybean, and sole cowpea) and soil amendments (Control, NPK, NPK+ biochar and sole biochar). Overall, the layout was a two-factor experiment in split – plot arranged in a randomized complete block design (RCBD) with cropping systems as main plot and soil amendments designated as sub-factor. The rates of each soil amendment varied with the crops as follows: 90-60-60 kg ha^-1^; 20-40-20 kg ha^-1^; 20-40-20 kg ha^-1^ of N, P_2_O_5_ and K_2_O for maize, soybean and cowpea respectively [12] and 5 t ha^-1^ of biochar [14, 15] while for the combination of the two amendments (inorganic and organic), 50 % NPK and 50% biochar were applied. The treatments were replicated three times. Each block comprised 16 plots with 16 treatments (4 × 4) and a bare plot, which served as erosion check. Each individual plot measured 12 m x 3 m separated from the subsequent one with aluzinc sheets fixed 0.5 m deep and 0.75 m high to avoid runoff contamination from the neighbouring plots. The field was divided into blocks based on the landscapes (slope) and three slope classes were defined: 3, 6 and 10% designated as slope 1, slope 2 and slope 3 respectively. Plate 1 describes the characteristics of an individual runoff plot. The observations were carried out in three consecutive growing seasons (2016-major, 2016-minor and 207-major) and the field was under natural rainfall regime.

### 2.2. Surface runoff measurement with tipping buckets

The runoff amount from the plots was collected at the base of each plot with the tipping bucket device (Plate 1). The tipping bucket device consisted basically of a collecting trough, tipping bucket and counter as described below:

#### Collecting trough

After the last row of crops, there was trapeze surface (covered by aluzinc sheets) to retain the first portion of runoff and sediments from the plot whilst the rest of water and the loads were passed through a mesh of 0.1 cm diameter for collection with the tipping bucket (Plate 2).

#### Tipping bucket devices and counter

After the mesh, the rest of water and its loads were passed through a channel of diameter 22.5cm, ending into a tipping bucket with two specific buckets (sides) with a known tipping volume for each (Plates 1). Once a bucket was filled with water or at the tipping volume, it tipped automatically and this was recorded from the counter fixed to the system. As a result, calibration of each of the devices was done each cropping season to confirm the tipping volumes. The volume of each bucket, obtained during the calibration process, was therefore used to calculate the total amount of runoff from each plot passing through the tipping bucket using equation (1) below.

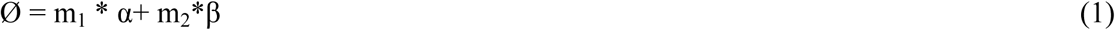

where. Ø (L). Total amount (volume) of runoff passed through the tipping buckets;
m_1_ (L) = tipping volume of the first bucket and m_2_ (L) = tipping volume of the second bucket. The tipping volume of each bucket was obtained at the tipping point during the calibration process carried out during each season;

α and β. Number of tipping times from the counter for the first and second buckets respectively.

The equation (2) was used to determine the total amount of runoff after subtracting the amount of water from the direct rainfall.

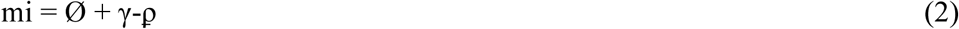

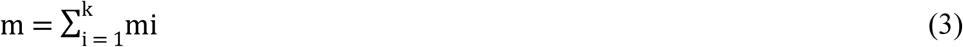

Where. mi (L). Total amount (volume) of runoff for an individual erosive rainfall;

> m (L). Total amount (volume) of runoff during k rainfall events;
>
> k . number of erosive rainfall events;
>
> γ (L). Volume of runoff collected from the small container (gallon) placed under the channel (sub-sample);
>
> Ø (L): Total amount of runoff from the tipping buckets and

ϼ (L): the volume of water from the direct rainfall collected in the collecting trough and which was determined using the equation (4).

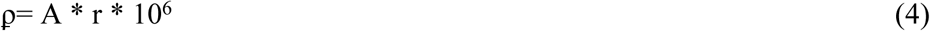

> Where: A (mm^2^) = area of the collecting trough which is trapezoidal;r (mm) = is the rainwater during each erosive storm event and
> 10^6^ is the conversion factor for water of mm^3^ into L).

### 2.3. Development of the new method for soil runoff measurement

The new method developed was based on mathematical equations described below:

#### 2.3.1. Procedures and theoretical approaches

By using the installed devices of tipping buckets, the total amount of runoff from each plot was collected through a uniform channel, with **N (cm)** as its diameter, and connected to the end of the plot (Plate 1). A line level was used for a good horizontality of the channel to ensure that the water was uniformly distributed to each space of N_i_ cm of the channel; and to be sure that the channel is not sloppy and that all the parts are on the same level of elevation.

A small tube with known diameter **n (cm)** was then fixed on the uniform channel to collect small portion of runoff into a small container (gallon) of **v** (L) as the volume.

The diameter of the channel; the small tube and the volume of the gallon for sub-sampling should depend on the rainfall characteristics of the zone. Knowing the maximum individual rainfall of the zone, this can help to decide on the sizes of the three parameters (N, n and v. This allowed for preventing any loss if the small container gets full before the sampling during a specific rainfall event. Mathematically, this is represented by equation (5) and this condition should be considered to avoid any flooding during the erosive rainfall. Thus, by using the principle of runoff coefficient, the container will never be full because the plot cannot lose the total amount of water received from the rainfall; even if the land is bare and very sloppy. The runoff coefficient depends on soil properties, soil moisture content, land cover, the slope and rainfall characteristics [16, 17] as well as the interaction between groundwater and surface water flows [18].

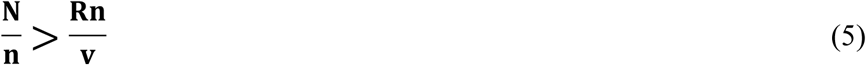

where: N (cm)= Diameter of the collecting channel;

> n (cm) = Diameter of the tube fixed on the channel;
>
> Rn (L) = Maximum amount of an individual rainfall of the study zone (this can be taken from the previous meteorological data during some years) that can be collected on a specific area;
>
> v (L) = volume of the small container for sub sampling the runoff.

#### 2.3.2. Runoff estimation or prediction

Following the above conditions and assumptions, the total amount of runoff for each erosive rainfall event (**pi**) and the total runoff during specific period of k rainfall events were determined by equations (6) and (7) respectively:

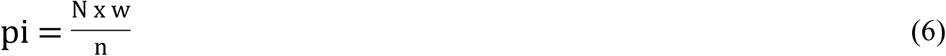

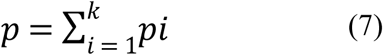

where: N (cm) = diameter of the collecting channel;

> n (cm) = diameter of the small tube fixed on the channel;
>
> w (L) = volume of runoff in the small container;
>
> pi (L)= individual predicted runoff for a specific erosive rainfall event;
>
> p (L)= total volume runoff predicted during a period of k erosive rainfall events and
>
> k = number of rainfall events during the study period.

### 2.4. Method quality evaluation and statistics analysis

Different statistic parameters were used for the quality assessment of the method developed. The goodness of fit between predicted and measured values was assessed using the statistical prediction errors. The coefficient of determination (R^2^), Nash –Sutcliffe efficiency (NSE), root mean square (RMSE), Root square ratio (RSR) were the parameters used to assess the quality of the method [19, 20]. The R^2^ and NSE allowed to access the predictive power of the model while RMSE indicates the error in model prediction [21]. The RSR incorporates the benefit of error index statistics and includes a scaling/normalization factor, so that the resulting statistics and values can apply to various constituents [22].

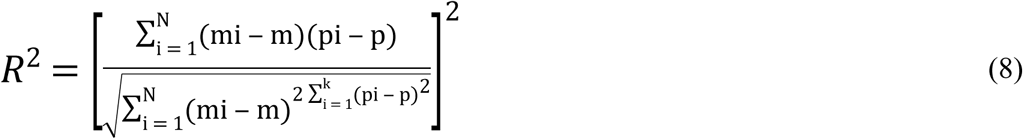

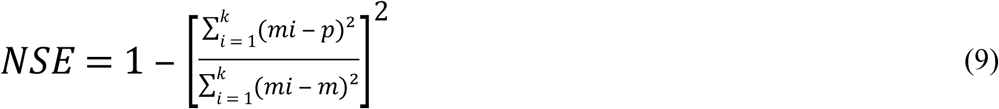

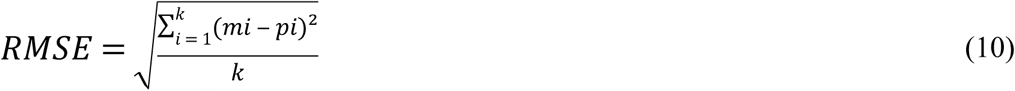

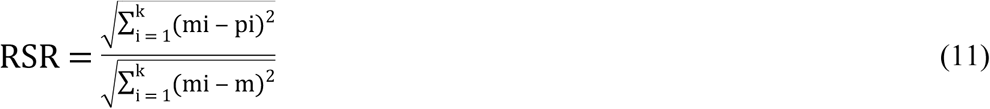

where: mi and pi = the measured and predicted values, respectively;

> m= the mean of measured values;
>
> p= the mean of predicted values and
>
> k= the number of observations (erosive rainfall events).

The data used for testing the models were measured from the 51 runoff plots in three consecutive cropping seasons: 2016-major, 2016-minor and 2017-major with 11, 9 and 13 erosive rainfall events, respectively. High number of observations allows for model accuracy [23]. Therefore, a total of 561, 459 and 663 direct observations were recorded during the three consecutive cropping seasons: 2016-major, 2016-minor and 2017-major seasons respectively for the model evaluation.

The different parameters used for the assessment were compared to their standards and ranges of acceptability as described by equations (8), (9), (10) and (11). For RMSE, lower values indicate better model agreement with predicted values. The coefficient of determination R^2^, the regression between measured and predicted values, ranges from 0 to 1, with higher values indicating better model prediction. NSE ranges between - and 1 and the values between 0.0 and 1 are generally considered as acceptable levels of performance. Negative values of NSE indicate that the mean of observed values is a better predictor than the simulated value, which indicates unacceptable performance of the model [22] . RSR varies from optimal value to 0, which indicates zero RMSE or residual variation and therefore perfect model simulation. Lower RSR values emphasize better model simulation performance. According to [22], the values are categorized as follow: 0.00 ≤ RSR ≤ 0.50; 0.50 < RSR ≤ 0.60; 0.60< RSR ≤ 0.70; RSR >0.70 for very good, good, satisfactory and unsatisfactory simulation, respectively.

For the effect of the different soil amendments and cropping systems on runoff, the analysis of variance (ANOVA) was perfumed using the least significant difference (LSD) method and the means separation at 5%. Prior to ANOVA, the data was checked for normality using residual plots in GENSTAT v. 12.

## 4. Results

### 3.1. Surface runoff variation under the different cropping systems and soil amendments

Under rain-fed cropping systems, rainwater is either used effectively by the plants or lost through different processes, especially runoff. The most important aspect is to increase the rainfall use efficiency by reducing unproductive water loss. As shown in Table 1, the different soil amendments and cropping systems affected rainfall water loss through runoff. Cropping systems significantly (P < 0.05) influenced runoff amounts during the three consecutive seasons. The sole cowpea reduced runoff more than the other cropping systems evaluated. Sole maize was the least tolerant to soil loss. The cropping systems reduced runoff in the order: Cw >S> M+S>M.

**Table 1.**
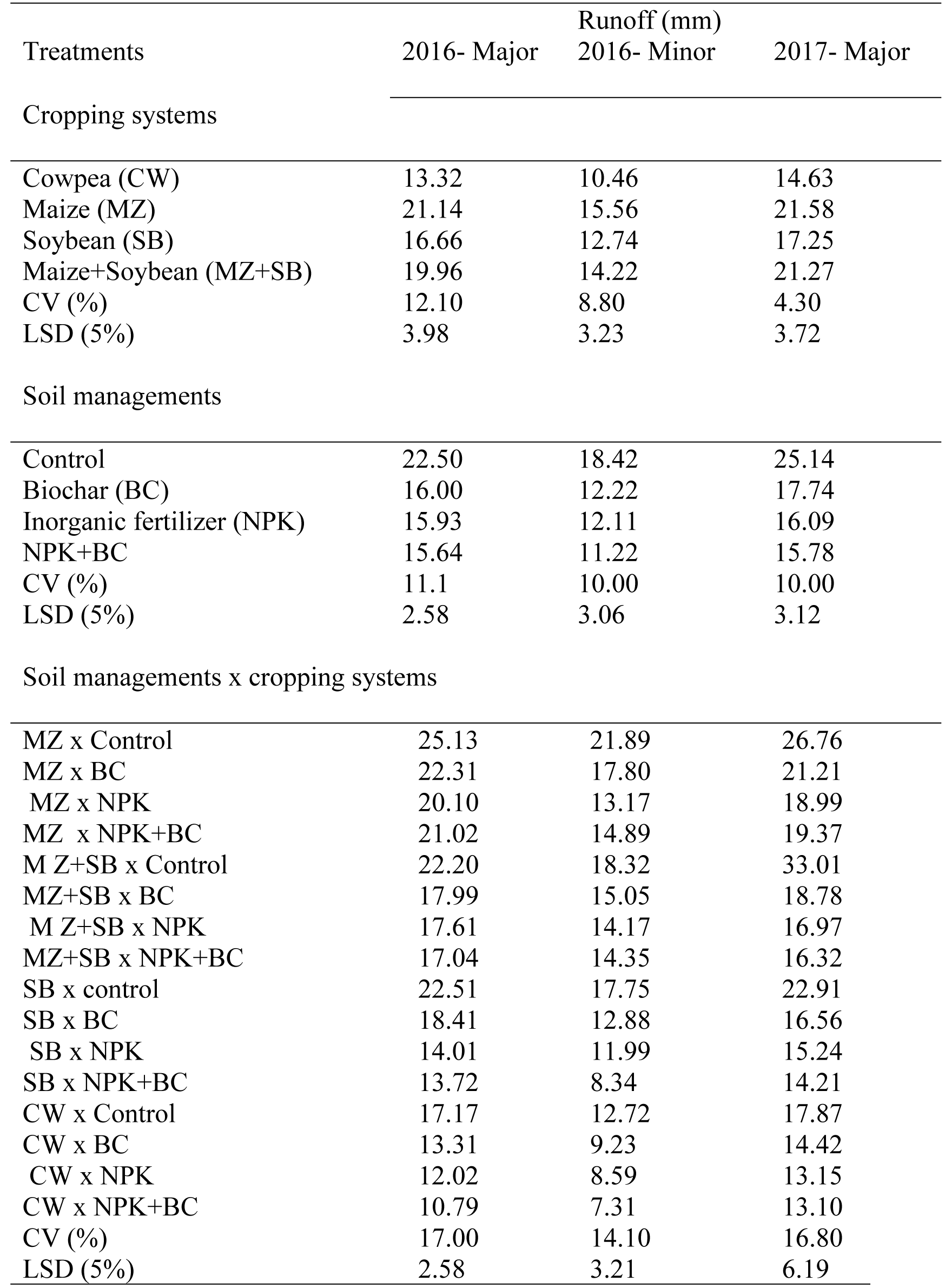
Effect of soil amendments, cropping systems and their interaction on runoff

For the soil management practices, the highest runoff loss was observed under the control ranging from 18.43 mm in the 2016 -minor season to 22.50 mm in the 2016-major season. The NPK + Biochar treatment reduced significantly soil runoff compared to the other amendments (Table 1).

The interaction effect between the two factors was significant (P< 0.05) with the highest runoff produced under sole maize without any amendment. Sole cowpea under NPK +Biochar consistently reduced runoff more than any other treatment combinations in all three seasons of cropping.

### 3.2. Characteristics of the new method for runoff estimation

The comparison between measured and predicted values for runoff is shown in Table (2). In general, all the factors of goodness presented excellent trends for a good model performance. The R^2^ and p-value between predicted and measured were R^2^ = 0.94 and p < 0.01; R^2^ = 0.94 and p < 0.01 and R^2^ = 0.89 and p < 0.01 in 2016–major, 2016-minor and 2017-major seasons respectively. The model showed good performance as the R^2^ values were close to 1 for all the three cropping seasons where 33 seasonal and cumulative erosive rainfall events were analyzed. The RMSE and RSR between measured and predicted runoff showed perfect thresholds with values of 2.67 and 0.40; 2.05 and 0.38 and 1.69 and 0.45 for the 2016-major, 2016-minor and 2017-major seasons, respectively. This showed that there was not much dispersion between measured and predicted values of runoff throughout the study period. For all the cropping seasons, NSE values ranged from 0.79 to 0.86, which qualified the prediction as excellent. The model showed good fit for runoff prediction through diagnostic plots of the linear model 1:1 (Figs 1a-1c).

**Table 2.** Performance indices between the predicted and measured runoff during different cropping seasons

| Index | 2016-major season | 2016-minor season | 2017-major season |
| --- | --- | --- | --- |
| $R^2$ | 0.94 | 0.94 | 0.89 |
| Slope | 0.56 | 0.66 | 0.56 |
| RMSE | 2.67 | 2.05 | 1.69 |
| RSR | 0.40 | 0.38 | 0.46 |
| NSE | 0.84 | 0.86 | 0.79 |
| p- value | <.001 | <.001 | <.001 |

**Figure 1 a.**
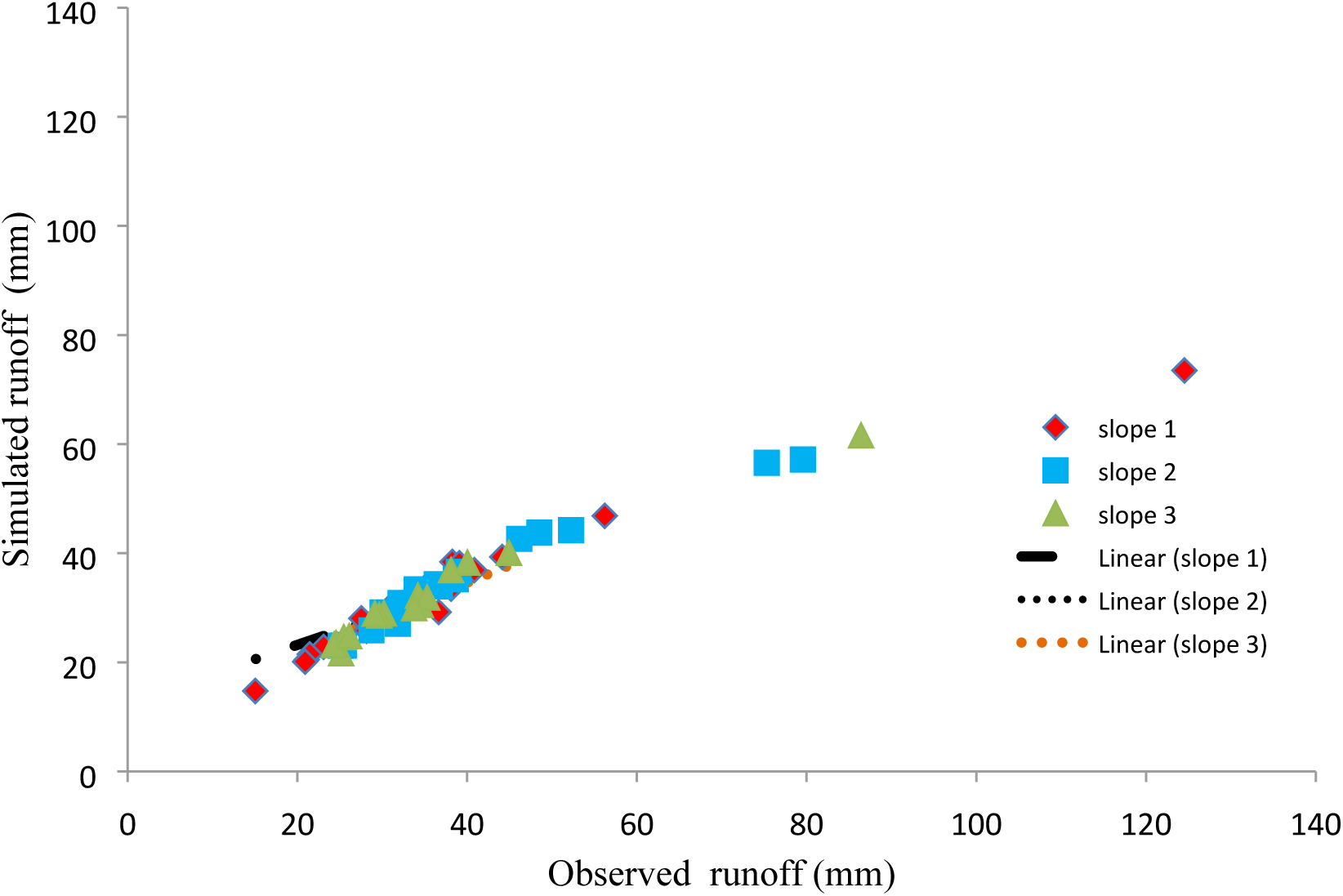
Effect of slope on model prediction under cropping systems and soil amendments during the 2016-major season

**Figure 1 b.**
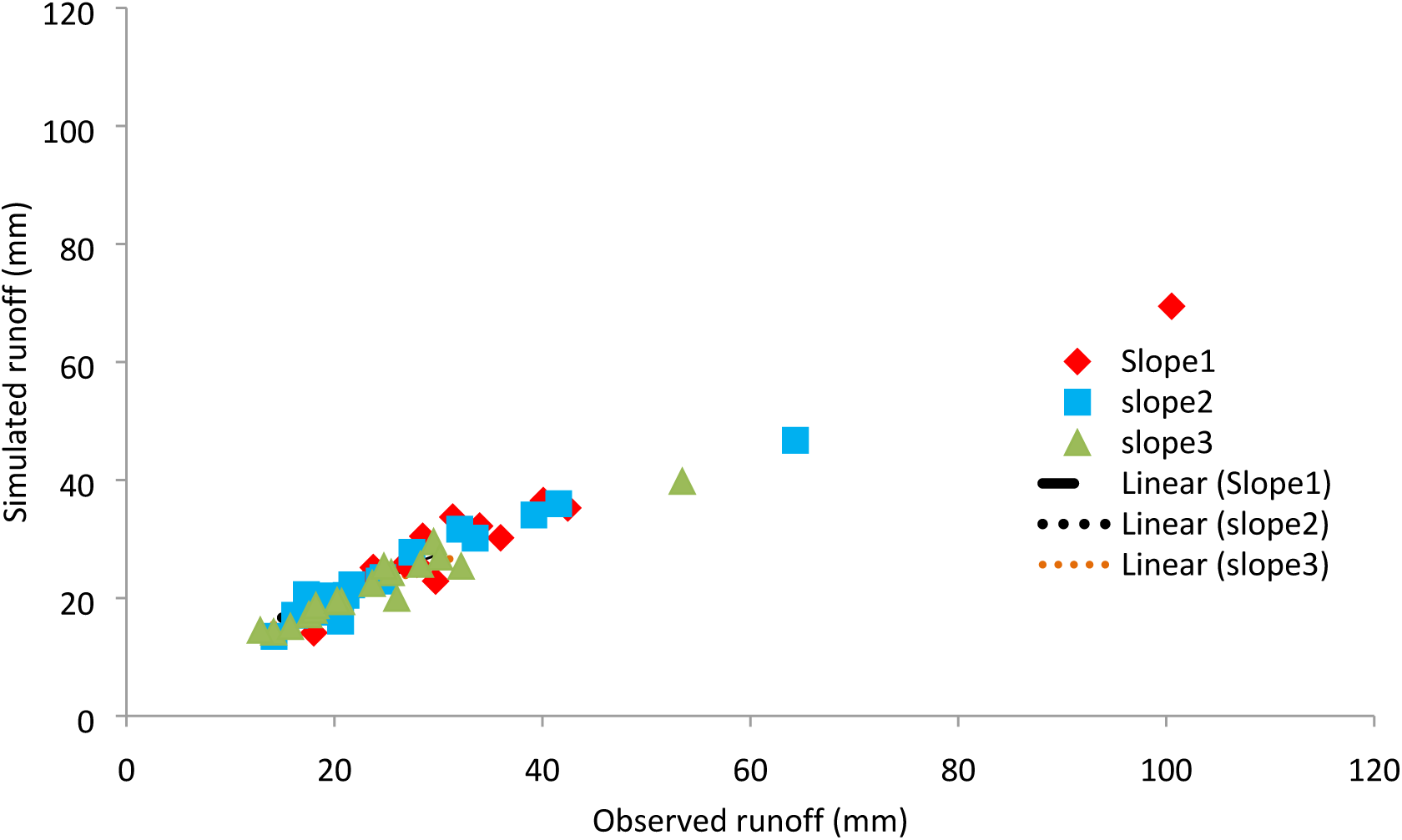
Effect of slope on the model prediction during the 2016-minor cropping season

**Figure. 1 c.**
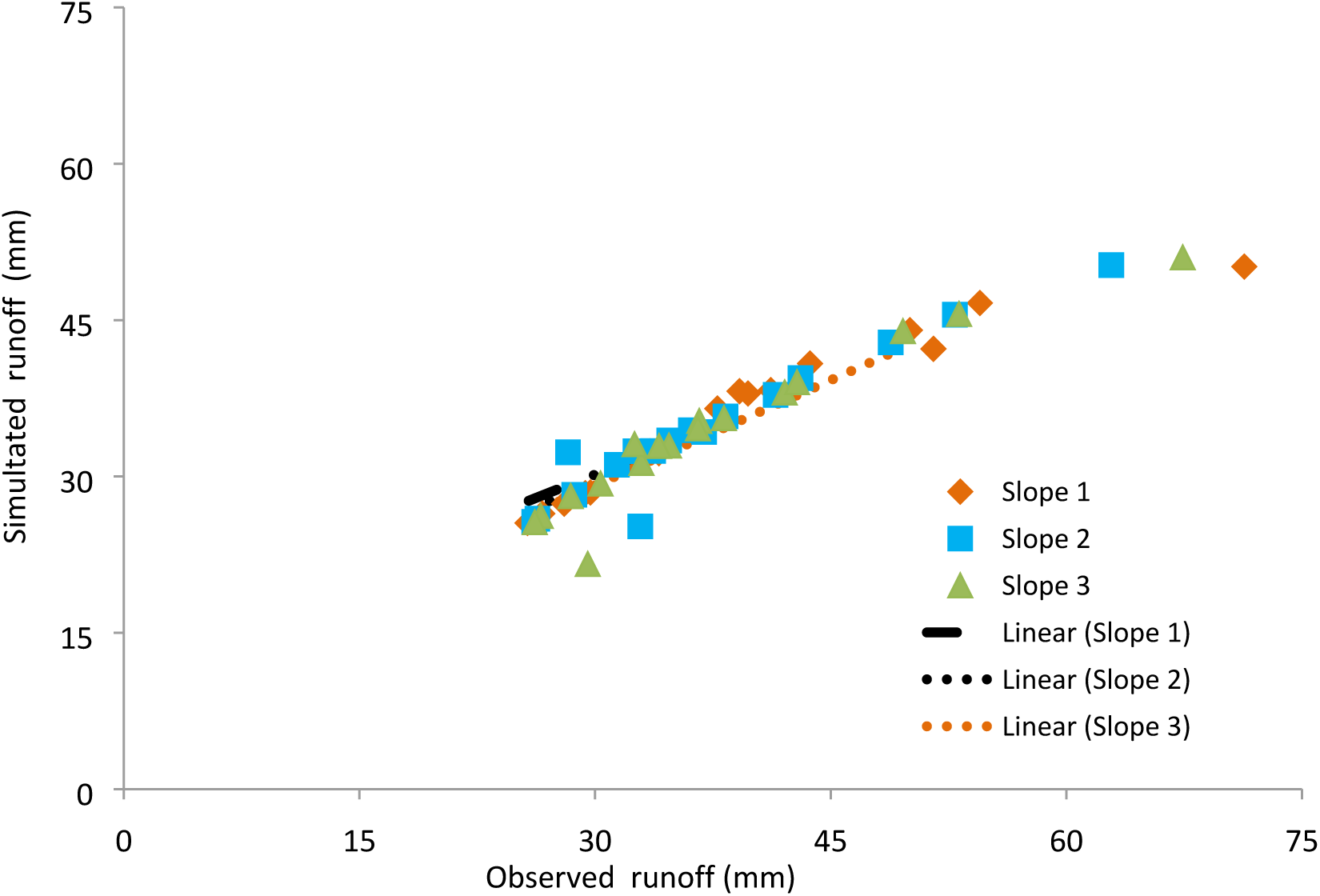
Effect of slope on the model prediction during the 2017-major cropping season

The accuracy of the runoff prediction under different slopes is presented in Figs. 1a-1c and the parameters (R^2^ and p-value) showed good performance and almost the same with the three slope classes (3, 6 and 10%). This confirmed that the current developed method could be applied to different landscapes based on slope steepness for soil erosion characterization

### 3.2. Sensitivity to different management and application of the model

The accuracy of the prediction is a function of the materials used for sub-sampling the runoff, which depend also on the climatic factor and the soil status as result of specific management practices and inherent properties. In Figs. 2b, 2d and 2f, the rainfall induced important amounts of runoff on poorly managed soils (bare plot).

From the equation (5), the variable N, n and v should be defined according to the rainfall characteristics (potential maximum daily rainfall amount) of the area for good accuracy of the simulation. The figures 2 a-2 f show good sensitivity of the model to predict the runoff under cropped and bare plots. The results showed good simulation as per the statistical parameters of goodness assessment (Table 2). All the figures without the bare plots (Figs 2 a, 2 c and 2 e) gave better accuracy of the prediction compared to the cropped plots mixed with the bare ones. Therefore, the bare plots with poorly managed soil, induced more runoff loss compared to the cropped land such that the estimation using the current method was poor for those three bare plots as marked with their respective peaks (^∗∗^) in Figs. 2b, 2d and 2f. The runoff was underestimated for the uncropped plots due to the high rate of the runoff generated and unsupported by the sampling tools. Under such circumstances, where high runoff occurs (Eq 5), the dimensions of the N, n and v should be adjusted to avoid losses due to overflow.

**Figure 2a.**
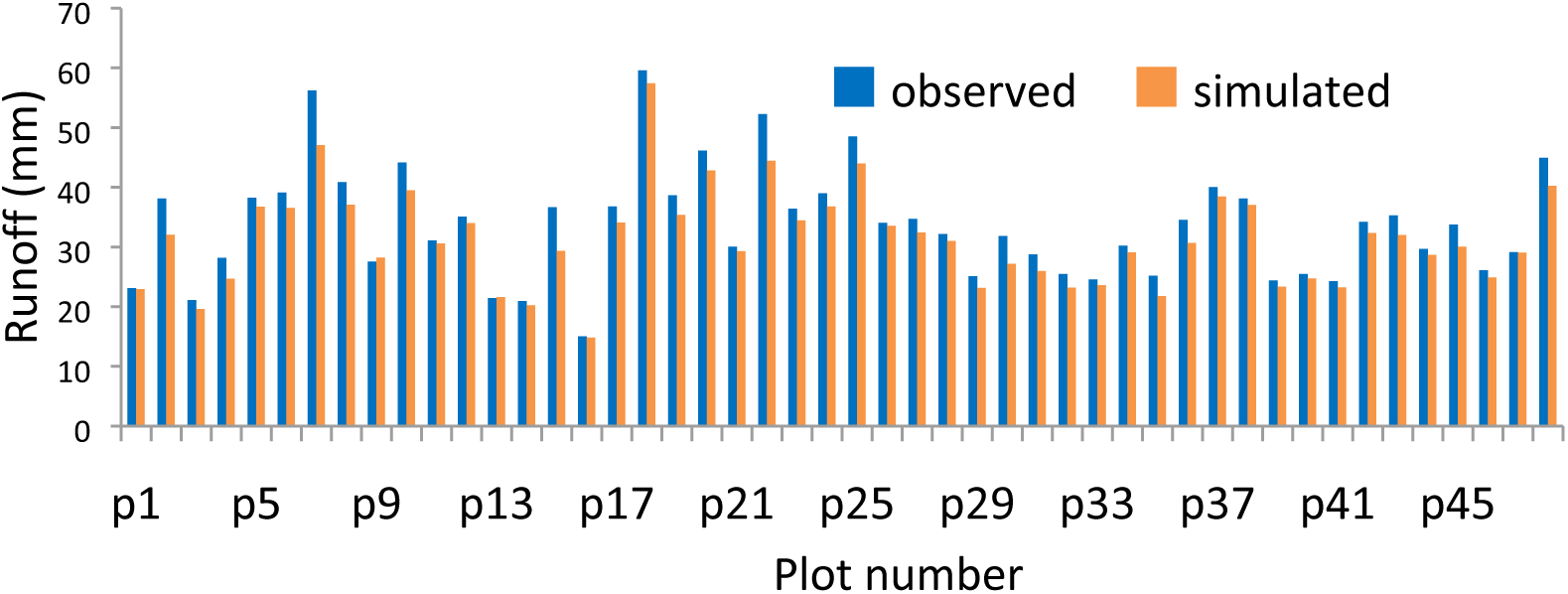
Runoff simulation and measurement sensitivity without bare plots during 2016-major cropping season

**Figure 2b.**
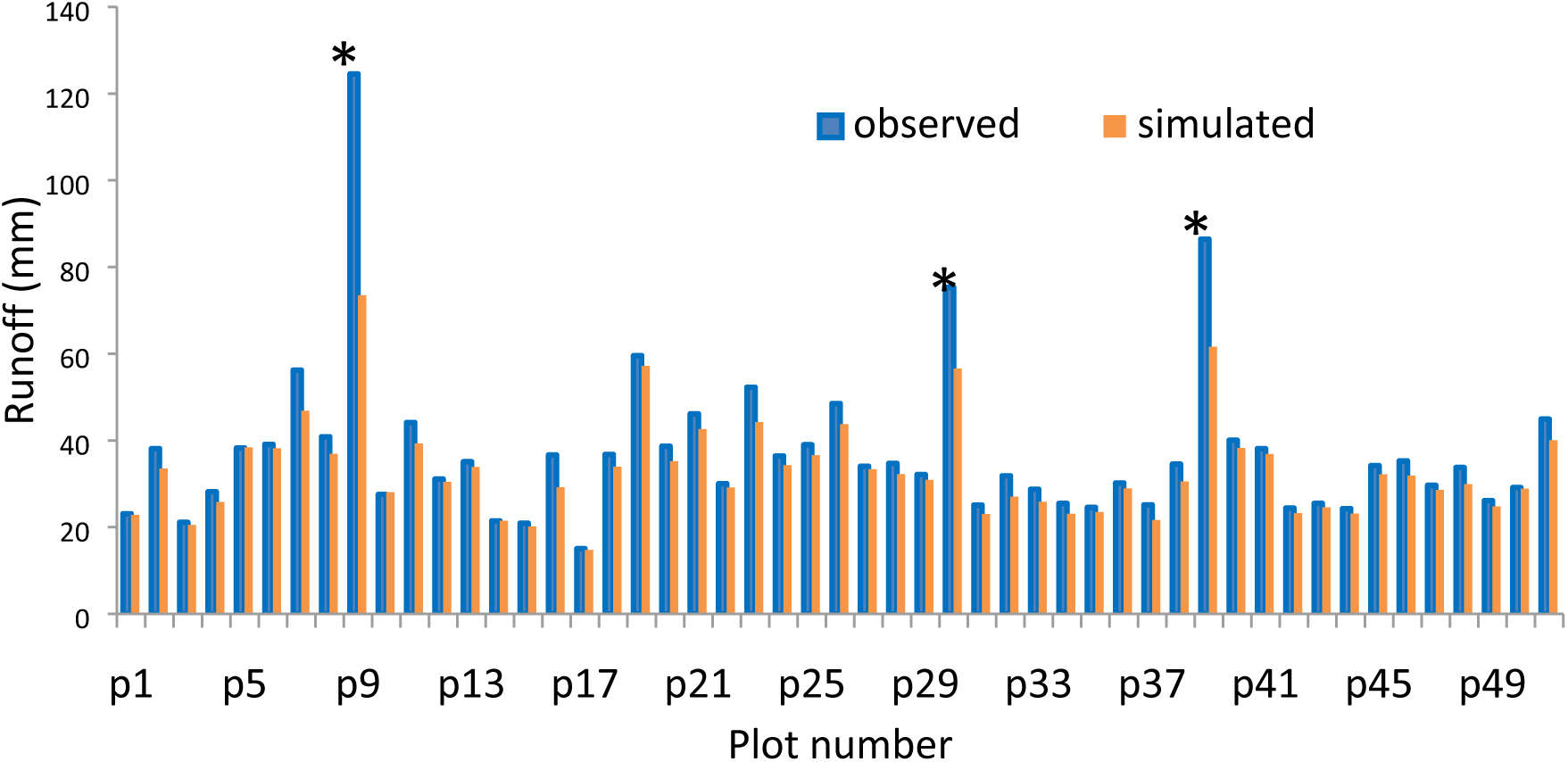
Runoff simulation and measurement sensitivity with bare plots during 2016-major cropping season. The ^∗∗^ on the three peaks of the bare plots show under-prediction when the flow is important compared to the cropped plots.

**Figure 2c.**
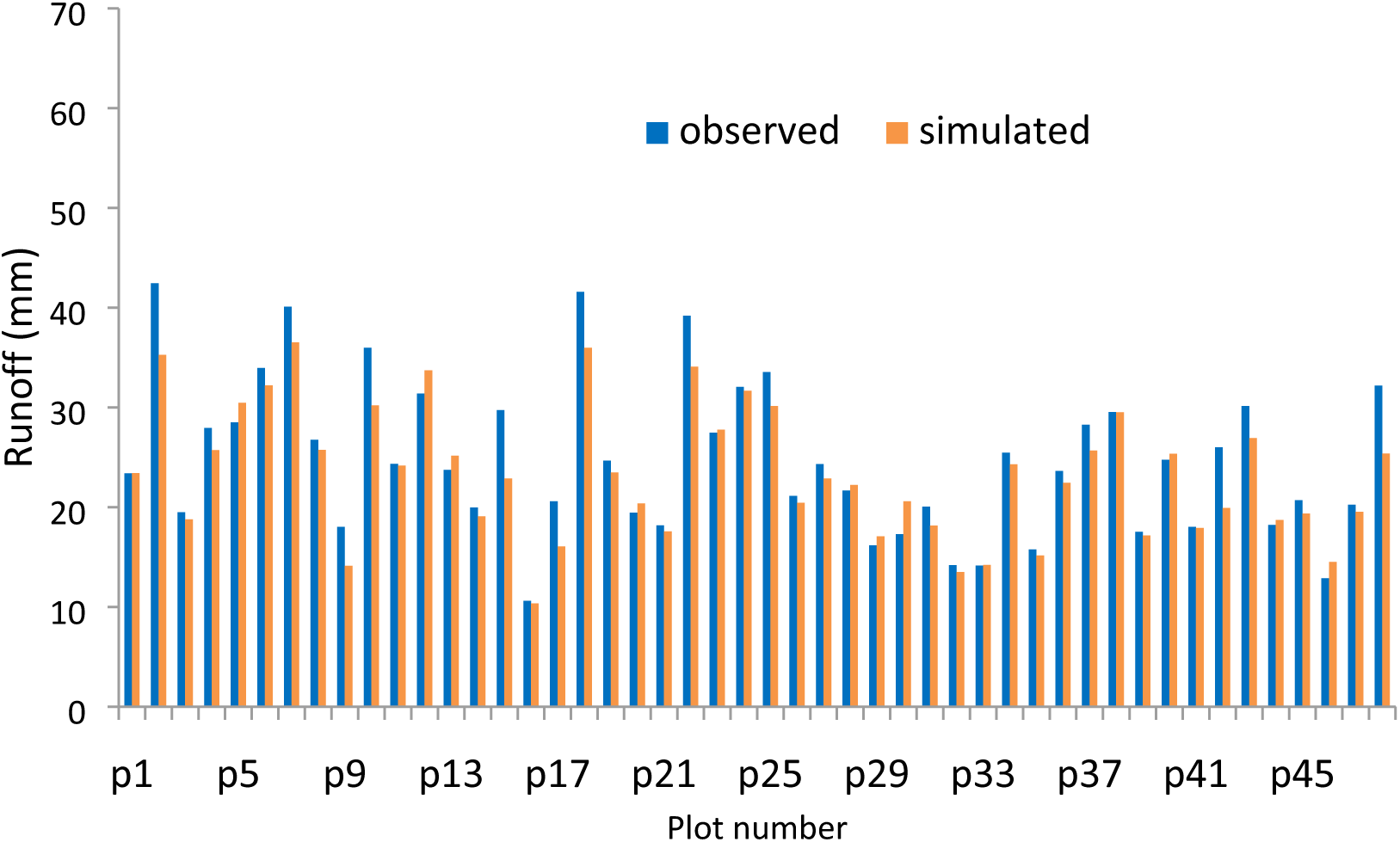
Runoff simulation and measurement sensitivity without bare plots during 2016-minor season

**Figure 2d.**
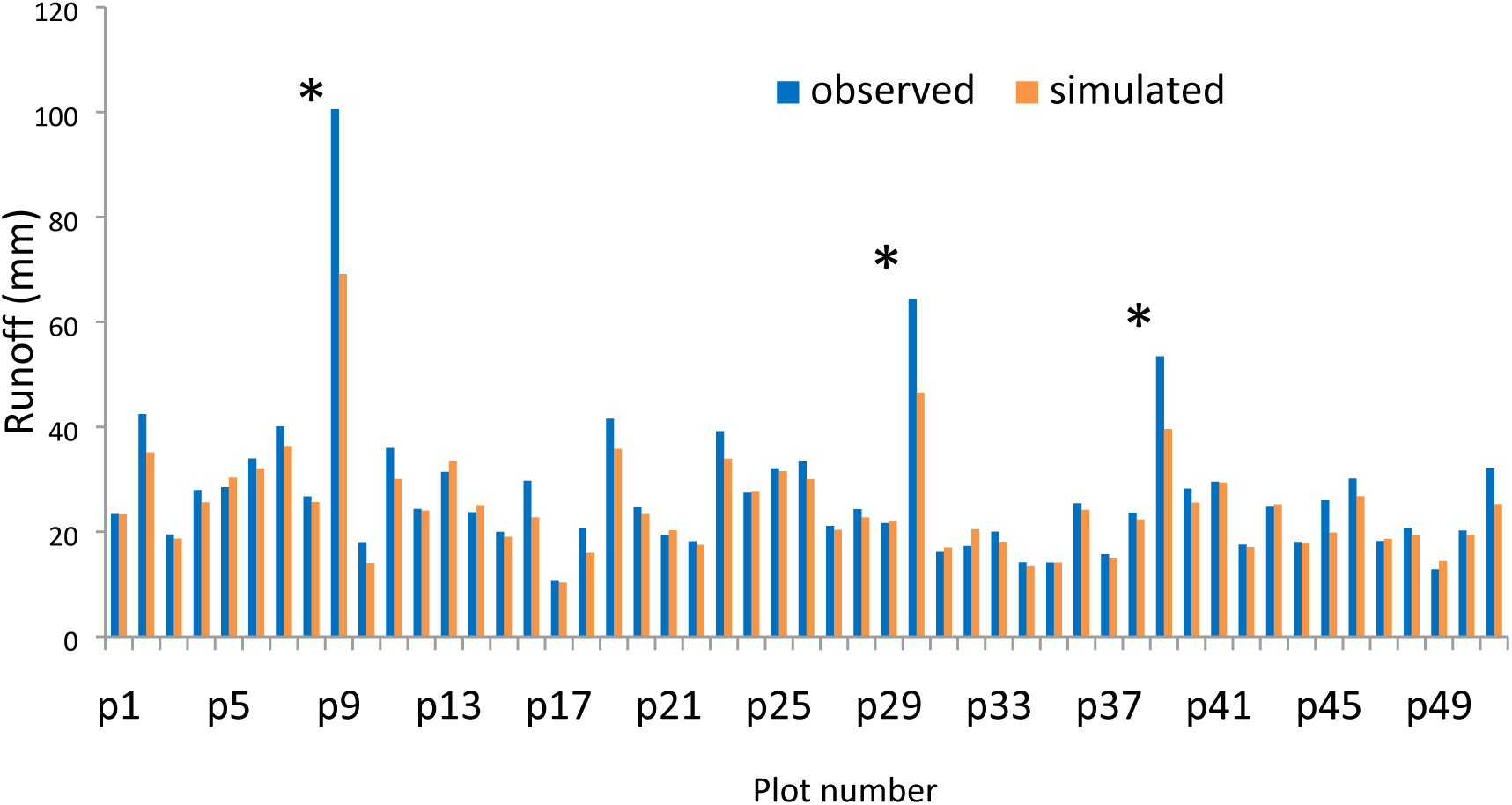
Runoff simulation and measurement sensitivity with bare plots during 2016-minor season. The ^∗∗^ on the three peaks of the bare plots show under-prediction when the flow is important compared to the cropped plots.

**Figure 2e.**
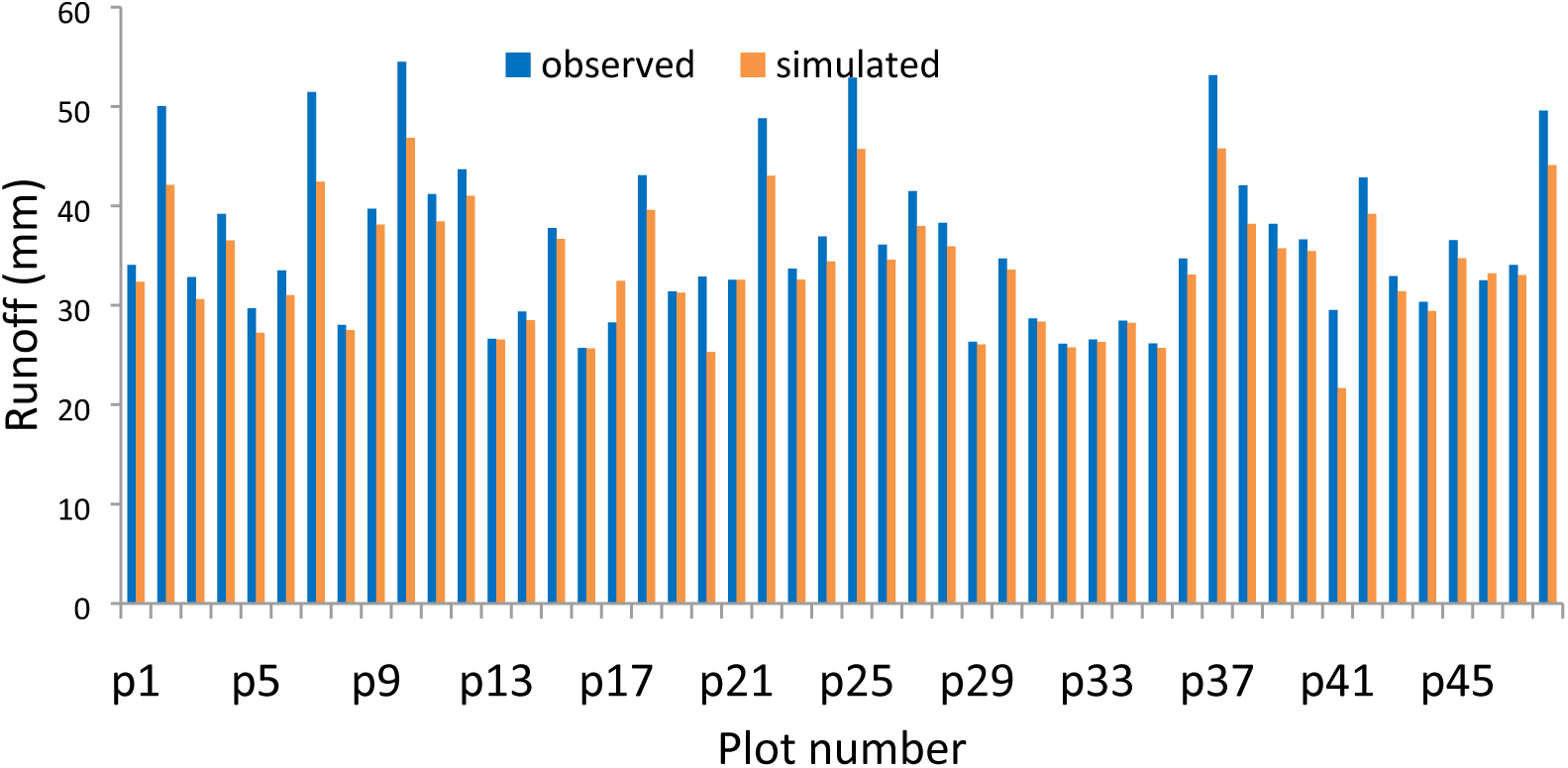
Runoff simulation and measurement sensitivity without bare plots during 2016-major cropping season.

**Figure 2f.**
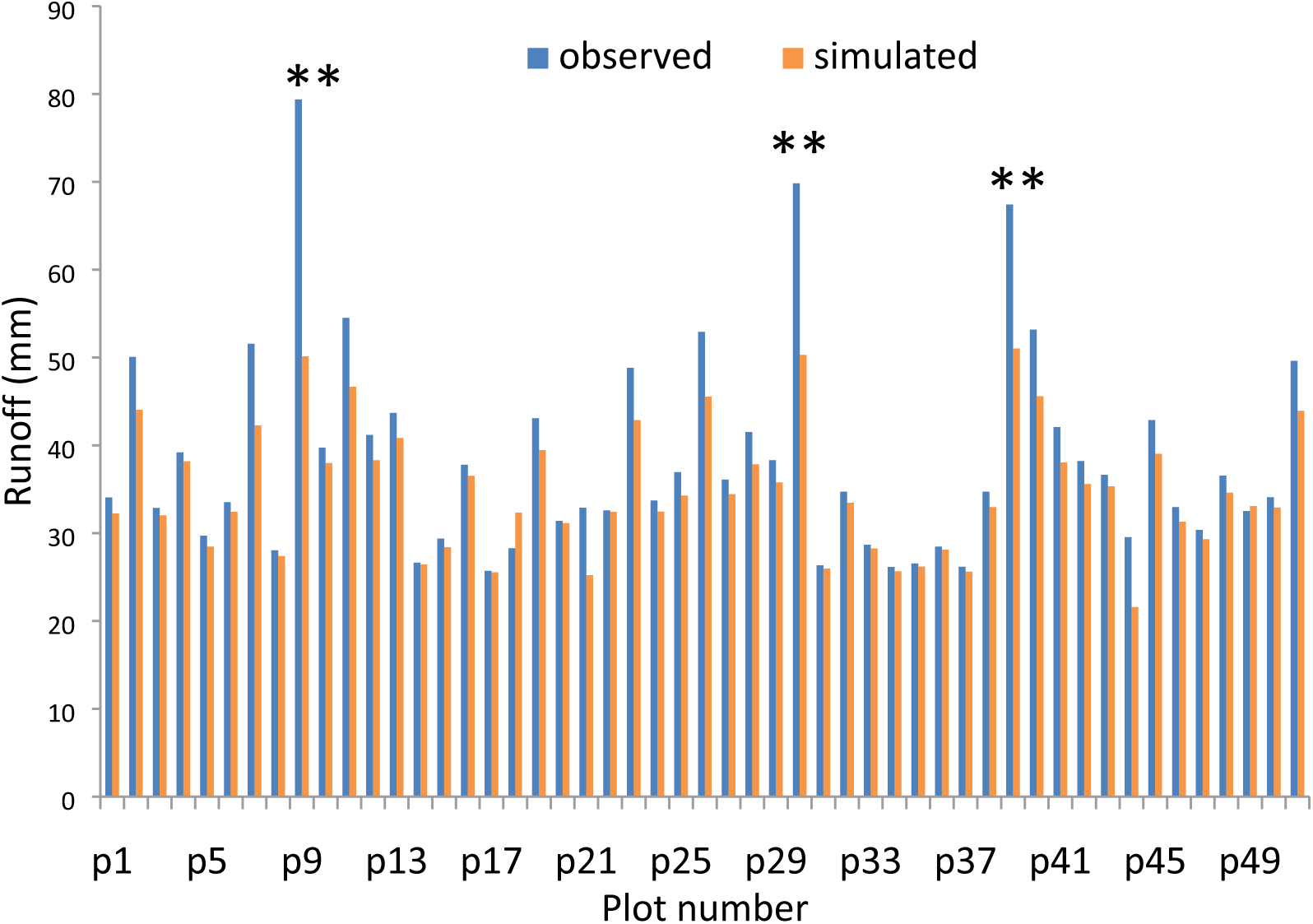
Runoff simulation and measurement sensitivity with bare plots during 2017-major season. The ^∗∗^ on the three peaks of the bare plots show under-prediction when the flow is important compared to the cropped plots

## 4.0 Discussion

### 4.1. Direct soil runoff measurement under the different cropping systems and soil amendments

Soil surface characteristics and soil management practices influence the fate of runoff generated under cropping systems. In this study, sole cowpea was more effective in reducing rainwater loss with the least amount of runoff followed by the sole soybean during the cropping seasons (Table 1). The decreased runoff observed under cowpea was due to its ability to provide better soil cover, which possibly reduced raindrop impact leading to increased rainwater infiltration and less runoff [24]. The role of cropping systems in reducing soil loss is based on provision of surface cover [25].

Soil nutrient management practices reduced also the amount of rainfall water lost through runoff during the three consecutive cropping seasons (Table 1) emphasizing the importance of plant growth, improved by sustainable nutrients supply on soil erosion management [26]. Even with its lower nutrients content compared to other soil organic amendments, biochar has positive effect on soil porosity and soil moisture storage [27] explaining the reduced runoff observed under this treatment in comparison with the control plots in this study (Table 1).

### 4.2. Accuracy assessment of the method

Several studies have used different and specific models to measure and predict soil erosion and runoff in assessing the impact of soil and crop management practices on soil and water management [28, 29, 30]. The selection of a specific model depends on the final objective of the study, the data required to run and calibrate it and the implicit uncertainty in interpreting the results obtained. However, the traditional physically-based, conceptual, and empirical or regression models developed have not been able to describe all processes involved due to insufficient knowledge and unrealistic data requirement. Thus, the application of most methods is limited to specific areas and studies.

The accuracies under the three slopes followed the same trends with good values of the coefficient of determination (R^2^ > 0.8) for each of the slope class as observed during the different cropping seasons (Figs 1 a-1 c). This confirmed the adaptability of the model under different types of landscapes based on slope as also was shown by the RSR values, which exhibited low variability among the different soil amendments. Thus, this gives a large applicability of the proposed approach for soil erosion characterization based on runoff determination within different landscape types. The adaptability of a model to different environments by keeping the same thresholds is one of the conditions to assess good model quality for soil erosion measurement [2].

Under the different cropping systems, a part from the bare plots, the prediction was accurate under the different soil management measures. The method satisfied the statistical thresholds of accuracy for runoff prediction as defined by [22] and the replicabiity under different soil management sytems based on the principles of [2].

### 4.2. Model application, advantage and limitation

The application of the actual model is based on the factors developed under equation 5 and the principle described by equations 6 and 7. Soil runoff quantified will therefore be used to assess the potential amount of soil and nutrients losses through erosion before suggesting sustainable practices for soil management and crop productivity improvement. Soil fertility restoration strategies will be based on the measured values of soil and nutrient loss to sustain agricultural production [31, 32]. The advantage of the proposed method is based on the following criteria: high accuracy under land management systems; applicability to different conditions including spatially varying soil and surface characteristics. As suggested by [2], a model with large conditions of adaptability and not specifically limited to certain situations are recommendable for soil erosion characterization under field and watershed scales. The current method is adapted and useful for soil erosion characterization on field scale basis. Contrary to other methods and models of soil runoff characterization where soil erosion is assessed after a long period of observation, as developed in different studies [eg. 2, 4, 5, 8, 33], the current method can assess the runoff for an an individual eosive storm. However, this new method of runoff assessment is limited with the design and size of the runoff plot to avoid any rainwater loss before sampling, as suggested by Plate 1.

**Plate 1.**
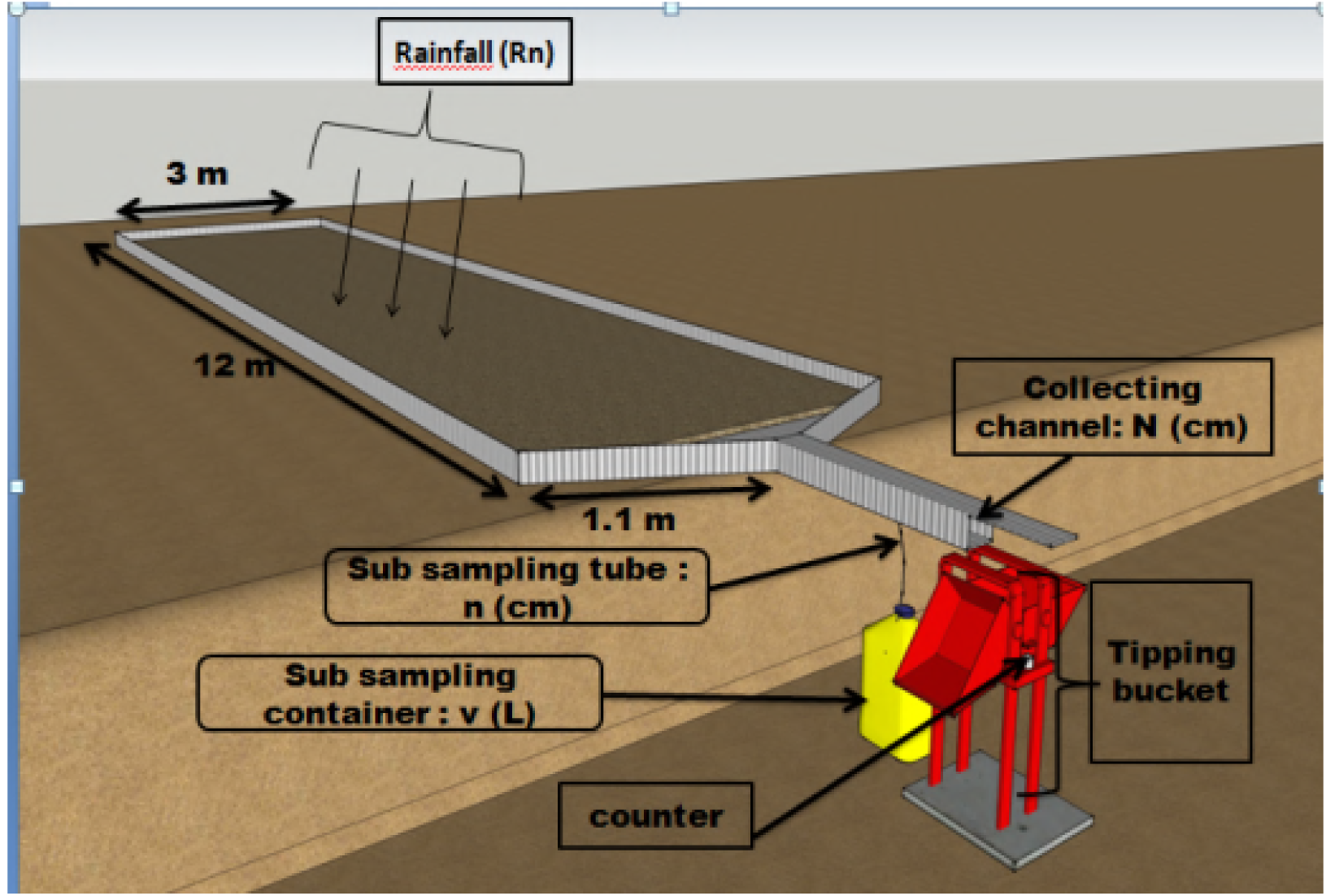
Layout of an individual runoff plot with the tipping bucket device for runoff and soil erosion assessment

**Plate 2.**
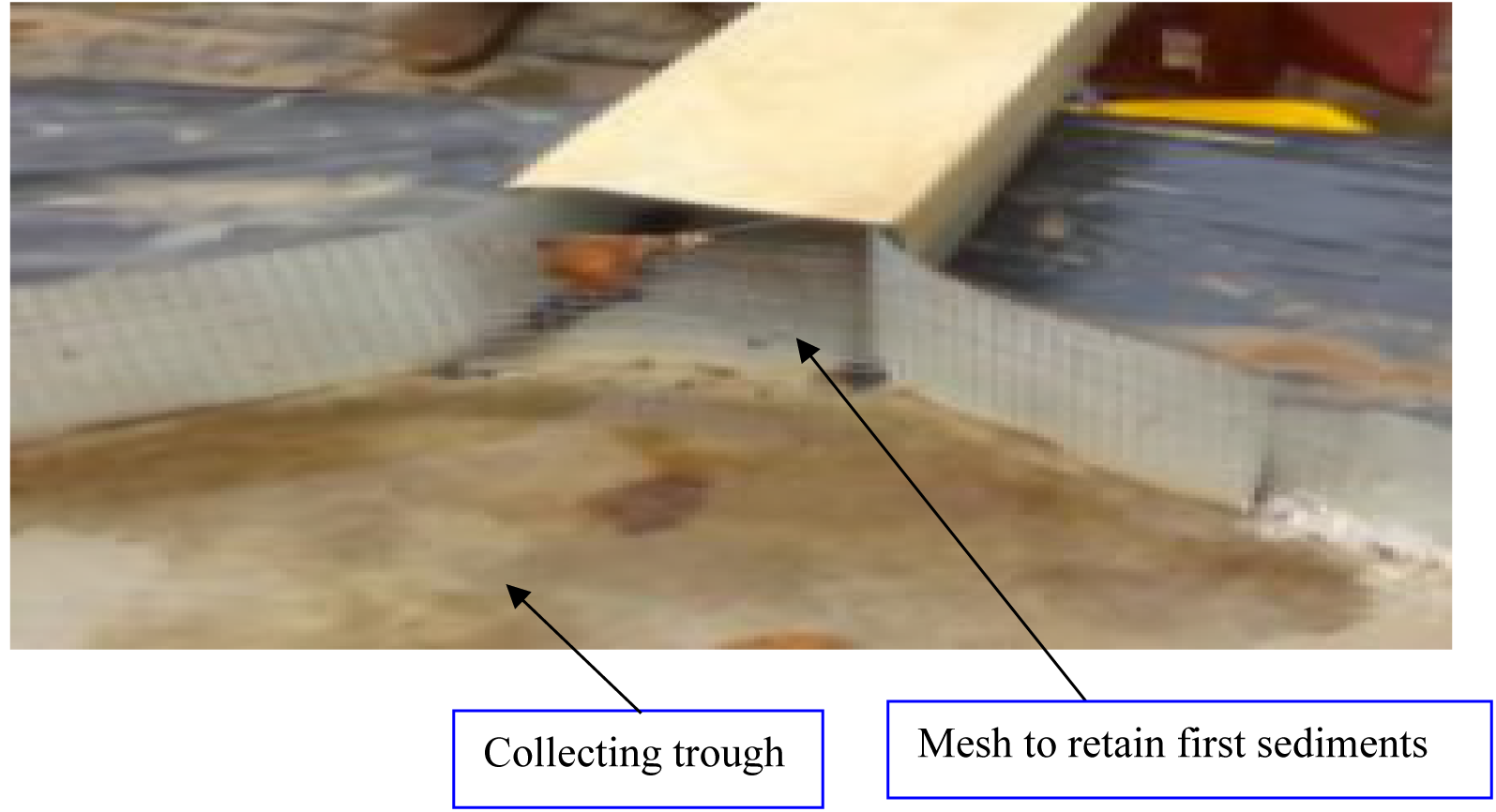
Collecting trough with aluzinc sheet at the end of each runoff plot and the mesh fixed between the channel and the collecting trough to retain the first portion of the runoff loads

## 5. Conclusion

The combined application of inorganic fertilizers and biochar was more effective under all the cropping systems in reducing runoff. Sole cowpea reduced runoff more than all cropping systems evaluated.

The developed model for soil runoff measurement was assessed using five statistical parameters of accuracy and goodness, which showed excellent thresholds and confirmed that the model performance for runoff prediction was accurate. All the five factors used for the assessment (p-values, R^2^, RMSE, NSE and RSR) gave excellence trends and as such the approach was qualified for soil erosion characterization. The model was assessed under different slope classes and showed good trends confirming its adaptability to different landscape types. Thus, this gives a new opportunity of soil erosion measurement under field conditions.

Despite the excellent predication of the method, the accuracy was poor for the plots with high rates of runoff (bare plots). Thus, the rainfall characteristics (runoff coefficient) of study regions in prospective works should be considered in fixing the characteristics of the collecting runoff. Although the statistical parameter based on RSR showed large adaptability trends, further test under different agro-ecological zones is recommended to assess the adaptability and the environmental effect on the accuracy of the method proposed.

## 6. Acknowledgment

This study was part of the PhD research of the lead author, supported by INTRA-ACP ACADEMIC MOBILITY project, in the Department of Crop and Soil Sciences of the Kwame Nkrumah University of Science and Technology (KNUST). Authors are also grateful to the research assistants particularly Mr Ayuba Salifu and the Anwomaso Research Station team for their supports during the study period.

